# Precision RNAi using synthetic shRNAmir target sites

**DOI:** 10.1101/2022.11.24.517802

**Authors:** Thomas Hoffmann, Alexandra Hörmann, Maja Corcokovic, Jakub Zmajkovic, Matthias Hinterndorfer, Jasko Salkanovic, Fiona Spreitzer, Anna Köferle, Katrin Gitschtaler, Alexandra Popa, Sarah Oberndorfer, Florian Andersch, Markus Schäfer, Michaela Fellner, Nicole Budano, Jan G. Ruppert, Paolo Chetta, Melanie Wurm, Johannes Zuber, Ralph A. Neumüller

## Abstract

Loss-of-function genetic tools are widely applied for validating therapeutic targets, but their utility remains limited by incomplete on- and uncontrolled off-target effects. We describe artificial RNA interference (ARTi) based on synthetic, ultra-potent, off-target-free shRNAs that enable efficient and inducible suppression of any gene upon introduction of a synthetic target sequence into non-coding transcript regions. ARTi establishes a scalable loss-of-function tool with full control over on- and off-target effects.

## Introduction

Drug development is guided by genetic loss-of-function (LOF) experiments that validate a therapeutic target, study its general and disease-specific functions, and thereby model and benchmark expected activities of inhibitory molecules. Applied genetic tools include RNAi, CRISPR/Cas9, and site-specific recombination technologies such as the Cre-Lox or Flp-FRT systems^1,2^. While these technologies have undoubtedly revolutionized genetic screening, target identification and validation, each method is associated with drawbacks that limit the usability for certain aspects of target validation. Specifically, RNAi is prone to off-target effects^3–5^ and insufficient knock-down levels, while CRISPR/Cas9-based methods are associated with off-target effects and incomplete LOF across cell populations. These limitations are particularly problematic for candidate targets in oncology, which should ideally be validated genetically in cancer cell lines and tumor models *in vivo*. In both cases, insufficient LOF or off-target effects can lead to far-reaching misconceptions about target suppression effects. Outgrowth of wild-type clones that retain gene function upon CRISPR- and recombination-based gene editing can result in transient phenotypes that complicate data interpretation. Similarly, insufficient knock-down or uncontrolled off-target effects induced by RNAi can lead to an unjustified de-prioritization or pursuit of candidate targets, respectively. Prior to initiating the time- and resource-intense process of drug development, more informative target validation assays would be highly desirable.

## Results

To develop such an assay system, we reasoned that instead of using gene-specific LOF triggers for every new candidate gene, the expression of any gene could be efficiently suppressed by engineering the target site of a pre-validated, highly potent, synthetic short-hairpin RNA (shRNA) into its exonic sequence (Figure 1A). Besides ensuring efficient target knockdown in a highly standardized manner, such an approach would also provide control over off-target activities through expressing the shRNA side-by-side in target-site engineered and wildtype cells. This approach leaves the genome engineering procedure, needed to integrate the pre-validated artificial RNA interference (ARTi) target site into a gene of interest, as the only potential source of off-target effects. As suitable RNAi system, we chose optimized micro-RNA embedded shRNAs (shRNAmirs) in the miR-E backbone^6^, which do not interfere with endogenous miRNA processing^7^ and can be expressed from tet-responsive elements and other Pol-II promoters in the 3’-UTR of fluorescent reporter genes, thus providing a versatile system for inducible RNAi^8^. To identify potent ARTi sequences with minimal off-target activity, we analyzed the nucleotide composition of shRNAs that reach exceptionally high performance scores in common shRNA design algorithms^9^ and of miRNA seed sequences with exceptionally low off-target scores according to siSPOTR^10^. By merging nucleotide biases identified in both analyses, we derived a 22-nt base composition matrix for the design of ARTi shRNAmirs (TTCGWWWNNAHHWWCATCCGGN; W = A/T, H = A/T/C; N = A/T/G/C) (Figure 1B; Figure 1 - figure supplement 1A,B). To further reduce possible off-target effects, we eliminated all shRNAs whose extended seed sequence (guide positions 2-14) had a perfect match in the human or mouse transcriptome and, finally, selected six top-scoring ARTi predictions for experimental validation.

**Figure 1:**
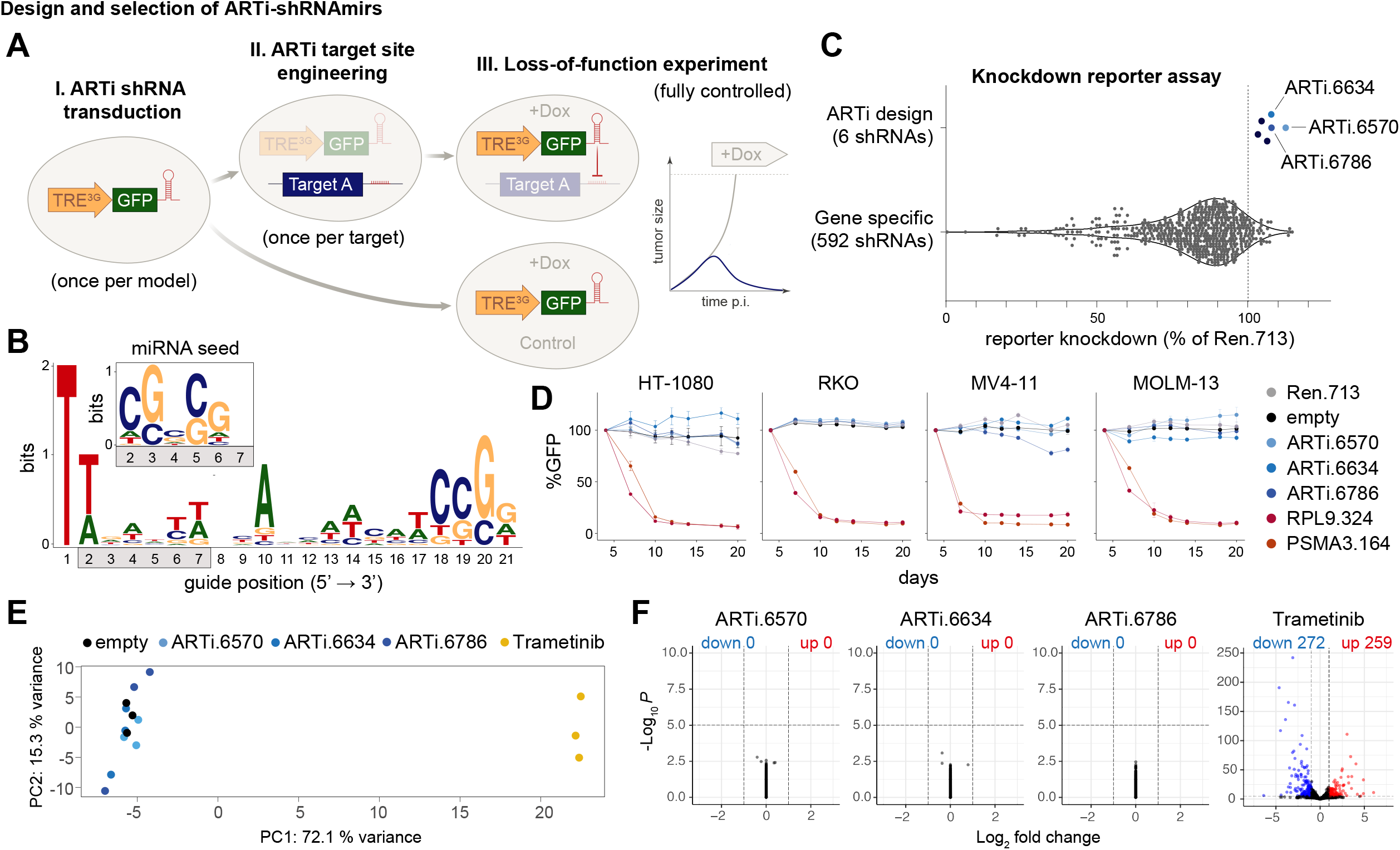
Design and selection ARTi-shRNAmirs. **A**, Schematic outline of the ARTi approach. **B**, Sequence logo (https://weblogo.berkeley.edu) displaying nucleotide position biases of 2161 shRNAs with exceptionally high DSIR scores (>105). Inlay depicts miRNA seed sequence biases. **C**, Reporter assay comparing gene-specific shRNAs to ARTi-shRNAmirs. **D**, Competitive proliferation assays in human cell lines after transduction with ARTi-shRNAmirs, neutral (shRen.713) and essential control shRNAs (shRPL9.324 or shPSMA3.164). **E**, Principal component (PC) analysis of gene expression profiling upon stable expression of indicated shRNAs or treatment with MEK inhibitor (trametinib) in RKO cells. **F**, Volcano plots visualizing de-regulated genes upon expression of indicated shRNAs and trametinib treatment in RKO cells, compared to empty vector control.

We tested these ARTi-shRNAmirs using an established knockdown reporter assay^6^ for their ability to suppress expression of a GFP transgene that harbors the respective target sites in its 3’-UTR. In all six cases, ARTi-shRNAmir matched or outperformed previously validated highly potent shRNAmirs targeting Renilla luciferase or PTEN^6^ (Figure 1 - figure supplement 1C). We compared these results to a panel of 592 gene-specific shRNAs that were tested using the same assay and found that ARTi-shRNAmirs ranked among top-performing shRNAmirs overall (Figure 1C). Next, we selected the three top-performing ARTi-shRNAmirs and evaluated possible off-target activities using competitive proliferation assays and transcriptome profiling. ARTi-shRNAmir expression had no effects on proliferation or survival in four human and three mouse cell lines (Figure 1D; Figure 1 - figure supplement 1D), with the exception ARTi.6634 and ARTi.6786, which induced a mild fitness defect in the murine leukemia cell line RN2 and MV4-11 respectively. In contrast to effects of a MEK inhibitor trametinib, which we included as a positive control, stable expression of ARTi-shRNAmirs had only marginal effects on the transcriptome in two commonly used human cell lines (Figure 1E,F; Figure 1 - figure supplement 1E,F) and no effect in proliferation assays (Figure 1 - figure supplement 1G), indicating that they do not trigger major off-target effects, even in the absence of their respective target site. Together, these studies established a set of highly potent, off-target-free ARTi-shRNAmirs, among which we selected ARTi.6570 (ARTi-shRNAmir) for further investigations.

To establish ARTi as a method for target validation, we performed ARTi-based LOF experiments in cancer cell lines and xenograft models for three prominent oncology targets: EGFR and KRAS, which act as driving oncogenes in various cancer types^11,12^, and STAG1, which has been identified as a synthetic lethal interaction with recurrent LOF mutations of STAG2^13–16^. To establish an ARTi-repressible version of oncogenic EGFR^del19^, we cloned ARTi-shRNAmir target sequences into an expression construct encoding EGFR^del19^ fused to dsRed (Figure 2A), which rendered Ba/F3 cells cytokine-independent and sensitive to EGFR inhibition (Figure 2 – figure supplement 2A). We introduced this construct into human EGFR^del19^-dependent PC-9 lung adenocarcinoma cells and subsequently knocked out the endogenous *EGFR* gene. The EGFR^del19^::V5::dsRed::ARTi transgene fully rescued the loss of endogenous EGFR^del19^, while doxycycline (dox)-induced expression of the ARTi-shRNAmir (ARTi.6570) strongly inhibited proliferation of PC-9 cells and triggered a near-complete suppression of the EGFR^del19^::V5::dsRed::ARTi protein (Figure 2B,C; Figure 2 – figure supplement 2B). Consistently, in RNA-sequencing we observed an almost complete drop of reads mapping to the codon-optimized *EGFR*^*del19*^*::V5::dsRed::ARTi* transgene upon dox-inducible expression of the ARTi-shRNAmir, as well as a downregulation of *DUSP6* and other canonical targets of RAF-MEK-ERK signaling, which acts as key effector pathway of EGFR^del19^ (Figure 2 – figure supplement 2C-E).

**Figure 2:**
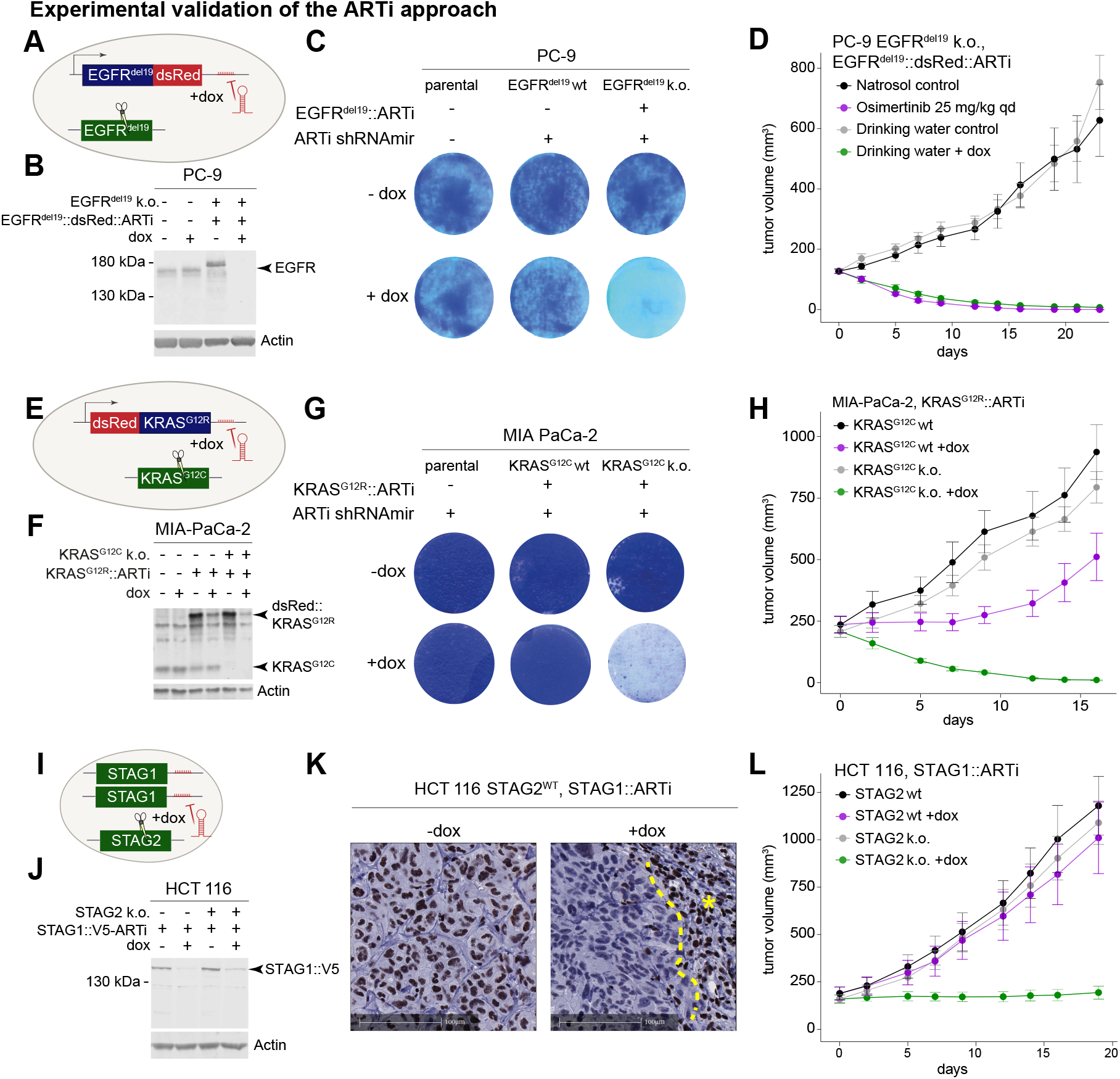
Experimental validation of the ARTi approach. **A**, Schematic of EGFR^del19^::V5::dsRed::ARTi engineering in PC-9 cells. Blue color denotes overexpressed ARTi variant. Green denotes endogenous gene. **B**, Western Blot demonstrating knock-down of EGFR^del19^::V5::dsRed::ARTi. Western Blot is a representative example of three independent biological repeat experiments. **C**, Proliferation assay and crystal violet staining of parental and engineered PC-9 cells in absence or presence of dox. Crystal violet staining is a representative example of two independent biological repeat experiments. **D**, *In vivo* experiment comparing dox-induced EGFR^del19^::V5::dsRed::ARTi knockdown to pharmacological EGFR^del19^ inhibition. Mean tumor volume and +/- s.e.m. is plotted for all *in vivo* experiments. **E**, Schematic of MIA PaCa-2 engineering. Blue color denotes overexpressed ARTi variant. Green denotes endogenous gene. **F**, Western blot for KRAS and Actin in indicated engineered MIA PaCa-2 cells in the presence and absence of dox. Western Blot is a representative example of two independent biological repeat experiments. **G**, Proliferation assay and crystal violet staining of parental and engineered MIA PaCa-2 cells in absence or presence of dox. Crystal violet staining is a representative example of two independent biological repeat experiments. **H**, Growth curves of tumors implanted with engineered MIA PaCa-2 cells in absence and presence of dox. In vivo **I**, Schematic of C-terminal endogenous tagging of STAG1. Green color denotes endogenous genes. **J**, Western Blot demonstrating knock-down of STAG1-ARTi. Western Blot is a representative example of three independent biological repeat experiments. **K**, Immunohistochemistry staining of STAG1 in engineered HCT 116 cells in absence and presence of dox. Asterisk marks an area of murine fibroblasts that serve as an internal positive control. **I**, Growth curves of tumors implanted with engineered HCT 116 cells in the absence and presence of dox.

To evaluate whether ARTi can recapitulate drug activities *in vivo*, we xenotransplanted *EGFR*^*del19*^*::V5::dsRed::ARTi* engineered PC-9 cells harboring the dox-inducible ARTi-shRNAmir and treated recipient mice upon tumor formation with either dox or the clinically approved EGFR inhibitor Osimertinib. Dox-induced expression of the ARTi-shRNAmir led to rapid and durable tumor regression that was indistinguishable from the effects of Osimertibib (Figure 2d) and not observed in parental PC-9 cells that lack the ARTi target site (Figure 2 – figure supplement 2F). We therefore conclude that ARTi-induced phenotypes are to be attributed to on-target effects that predict the activity of advanced small-molecule inhibitors *in vivo*.

In a second study, we used ARTi to investigate KRAS^G12R^, an oncogenic KRAS variant with no available *in-vivo* compatible inhibitors. To establish a suitable KRAS^G12R^-driven model, we engineered a *dsRed::KRAS*^*G12R*^*-ARTi* transgene and the dox-inducible ARTi-shRNAmir into KRAS^G12C^-dependent MIA PaCa-2 pancreatic adenocarcinoma cells and subsequently knocked out endogenous *KRAS* alleles (Figure 2E; Figure 2 – figure supplement 3A). As expected, switching the driving oncogene from KRAS^G12C^ to KRAS^G12R^ rendered MIA PaCa-2 cells resistant to the KRAS^G12C^ inhibitor AMG-510^17–19^ (Figure 2 – figure supplement 3B). ARTi-shRNAmir induction led to strong suppression of dsRed::KRAS^G12R^ expression at the mRNA and protein level (Figure 2F; Figure 2 – figure supplement 3C,D) and marked anti-proliferative effects (Figure 2G). In xenograft experiments, ARTi-mediated suppression of KRAS^G12R^ led to full tumor regression in the absence of KRAS^G12C^ (Figure 2H), while tumors harboring both oncogenic KRAS alleles only displayed a delay in tumor progression, suggesting that both oncogenes contribute to tumor growth *in vivo*.

To evaluate STAG1 as synthetic-lethal dependency that is relevant in wide range of STAG2-deficient cancers (Figure 2 – figure supplement 4A)^13–16^, we engineered an isogenic pair of STAG2-wildtype and - deficient HCT 116 colon carcinoma cells and homozygously inserted ARTi target sites (besides an AID::V5 tag; explained in the materials and methods section) into the 3’-UTR of endogenous *STAG1* (Figure 2I). Western blotting confirmed the knockout of *STAG2*, the insertion of the targeting cassette into the *STAG1* locus, and potent suppression of STAG1 following dox-induced ARTi-shRNAmir expression (Figure 2J,K; Figure 2 – figure supplement 4B). Suppression of STAG1 in STAG2-deficient HCT 116 cells impaired their proliferation *in vitro* (Figure 2 – figure supplement 4C) and the progression of xenografted tumors *in vivo* (Figure 2L), while dox-induced expression of the ARTi-shRNAmir had no anti-proliferative effects in the presence of STAG2.

## Discussion

Together, these studies establish ARTi as a versatile and precise LOF method for target validation in oncology and beyond. Instead of designing gene-specific LOF reagents that remain prone to off-target effects, ARTi involves a simple, highly standardized experimental procedure that provides full control over on- and off-target effects and can be applied to any coding and non-coding gene. In cells that are pre-engineered to express the dox-inducible ARTi-shRNAmir, candidate genes can be converted into ARTi target genes and subsequently evaluated head-to-head in a highly standardized manner, either through knocking in ARTi target sites into endogenous loci or through knock-out rescue approaches. Placement of ARTi target sites in non-coding transcript regions leaves the endogenous target protein unaltered, which is a key advantage over chemical-genetic LOF methods relying on the introduction of degron tags^20,21^. Targeting endogenously engineered ARTi target sites through tetracycline (Tet)-inducible ARTi-shRNAmir expression enables inducible, titratable, and likely reversible (not tested in this study) LOF perturbations of candidate genes without altering their endogenous transcriptional regulation, which is not possible using conventional Tet-expression systems. In principle, by engineering multiple target sites in the same cell, ARTi enables combinatorial LOF perturbations, e.g. for modeling synergistic target interactions, without multiplying the risk for off-target effects. Although ARTi requires the design of gene-specific sgRNAs and genome engineering steps that can be non-trivial, the method enables researchers to reach well-interpretable, comparable, and unambiguous experimental results that can guide larger investments in targets of high medical interest. Beyond providing a scalable method for early target validation, we foresee that ARTi can be used to establish on-target benchmark phenotypes for guiding the development and optimization of inhibitory molecules.

## Figure Legends

**Figure 1 – figure supplement 1: Design and selection ARTi-shRNAmirs. A**, Sequence logo displaying nucleotide position biases of seed sequences (guide position 2-8) with the lowest off-target score in siSPOTR analysis (top 1%). **B**, Sequence logo displaying position biases in sequences from a that harbor a T in guide position 2. **C**, Knockdown efficiency of ARTi-shRNAmirs. Flow cytometric quantification of GFP knockdown efficiency in immortalized MEFs two days after transduction with indicated ARTi-shRNAmir or control shRNAs. Percent knockdown is normalized to shRen.713 control. shRen.660 served as a neutral control whose target site was not included in the reporter. Red asterisks indicate ARTi-shRNAmirs that were selected for follow-up studies. **D**, Toxicity of ARTi-shRNAmirs. Competitive proliferation assays of three murine cell lines after transduction with ARTi-shRNAmirs, shRen.713 or shMyc.1834 control, showing the relative fraction of shRNA-expressing cells at indicated time points following the initial measurement (day 4 after shRNA transduction). **E**, Principal component analysis of gene expression profiling in HT-1080 cells. Respective shRNAmirs and treatments are indicated in the respective colors. X-axis: principal component 1; Y-axis: principal component 2. **F**, Volcano plots visualizing de-regulated genes in HT 1080 cells. All shRNAmirs and treatments were tested against the empty vector control. X-axis: - log_10_(p-value); Y-axis: log_2_ fold change. **G**, Cell growth assay for ARTi-shRNAmir transduced cells and their parental controls in the presence and absence of dox.

**Figure 2 – figure supplement 1: Validation of ARTi *in vitro* and *in vivo* - EGFR. A**, Functional validation of the EGFR^del19^ construct in Ba/F3 cells. Provided are GI_50_ values (nM) for indicated conditions and compounds (parental cells IL-3 dependent; other cells dependent on transgenes (no IL-3)). Numbers represent GI_50_ value calculated from three technical repeats. Three biological repeat experiments were conducted, and one representative experiment is shown in panel A. **B**, Western Blot analysis confirming expression of EGFR^del19^::V5::dsRed::ARTi constructs, knockout of endogenous EGFR and dox-induced knockdown or EGFR^del19^::V5::dsRed::ARTi. Western Blot is a representative example of three independent biological repeat experiments. **C**, Expression levels (y-axis: normalized counts) of the EGFR^del19^::V5::dsRed::ARTi construct in parental and engineered PC-9 cells after 4 and 8 days of dox treatment or no treatment. Individual datapoints of three biological replicates overlay the boxplots. **D**, Expression levels (y-axis: normalized counts) of the MAPK target gene *DUSP6* in parental and engineered PC-9 cells after 4 and 8 days of dox treatment or no treatment. Individual datapoints of three biological replicates overlay the boxplots. **E**, Heatmap visualizing the gene expression changes in MAPK pathway target and ERBB genes in engineered cells in the presence and absence of dox. **F**, *In vivo* experiment comparing dox-induced expression of the ARTi-shRNAmir to drinking water control in PC-9 parental cells. Lines connect mean tumor volume data and +/- s.e.m..

**Figure 2 – figure supplement 2: Validation of ARTi *in vitro* and *in vivo* - KRAS. A**, Schematic of MIA PaCa-2 genome engineering. **B**, AMG-510 GI_50_ values in MIA PaCa-2 parental and engineered cells. Numbers represent GI_50_ value calculated from three technical repeats. Three biological repeat experiments were conducted, and one representative experiment is shown in panel B. **C**, Expression levels (y-axis: normalized counts) of the dsRed::KRAS^G12R^ transgene in parental and engineered MIA PaCa-2 cells after 4 and 8 days of dox treatment or no treatment. Individual datapoints of three biological replicates overlay the boxplots. **D**, Western blot for KRAS (green) and Actin (magenta) (upper plot) and dsRed (green) and Actin (magenta) (bottom plot) for indicated engineered MIA PaCa-2 cells in the presence and absence of dox. Western Blot is a representative example of three independent biological repeat experiments.

**Figure 2 – figure supplement 3: Validation of ARTi *in vitro* and *in vivo* – STAG1. A**, Schematic of synthetic lethal interaction between STAG1 and STAG2. Cells survive loss of either paralog but are incapable of growing upon combined loss of STAG1 and STAG2. **B**, Western blot confirmation of STAG2 knockout and ARTi-shRNAmir induced knockdown of endogenous STAG1::V5::ARTi. Western Blot is a representative example of three independent biological repeat experiments. **C**, Proliferation assay of ARTi engineered HCT 116 cells *in vitro*, visualized by crystal violet staining following a 9-day dox or control treatment. Staining is a representative example of three independent biological repeat experiments. **D**, Quantification of nuclear Stag1 level (Figure 2k) using engineered HCT 116 cells in an in vivo xenotransplantation experiment the control group (-dox) and doxycycline (+dox) treated group.

**Figure 2 – source data 1**. Original blots for Figure 2B and Figure2 - figure supplement 1B.

**Figure 2 – source data 2**. Original blots for Figure 2F.

**Figure 2 – source data 3**. Original blots for Figure 2J.

**Figure 2 – figure supplement 2 – source data 1**. Original blots for Figure2 - figure supplement 2D.

**Figure 2 – figure supplement 3 – source data 1**. Original blots for Figure2 - figure supplement 3B.

## Material and Methods

### Design and cloning of ARTi shRNAs

To design pairs of artificial shRNAs and matching target sites that trigger effective and selective target suppression with minimal off-target effects, nucleotide composition of shRNAs that reach exceptionally high performance scores in a well-established siRNA prediction tool (DSIR=Designer of Small Interfering RNA^9^) and contain no A or T in guide position 20 to eliminate shRNAmirs that produce RISC-loadable small RNAs from the passenger strand were analyzed. To establish criteria for ARTi design, siRNA predictions for the human and mouse genome were retrieved from DSIR^9^ using default parameters. Top scoring predictions (DSIR score > 105) harboring G or C in position 20 were analyzed for nucleotide biases, which were used to define basic design criteria at the 5’-end the 3’-half. Next, possible off-targets were minimized using siSPOTR^10^, an siRNA-based prediction tool that assesses off-target potential of different siRNA seed sequences (guide positions 2-8) in the human and mouse genome. Seed sequences with the lowest predicted off-target activity (top 1%) showed biases for C and/or G in guide positions 2-6, but 23% contained a T at the 5’ end of the seed sequence that is required in our design (Fig. 1b). Among these, we observed particularly strong biases for CG in the following two positions (Fig. 1b), which were found in 73% of all seed sequences harboring a 5’ T. Overall, 17% of all top-scoring seed sequences harbored TCG at their 5’ end, making it the second most common triplet (after CGC, which was found in 20%). Based on these analyses, we fixed the first 4 nucleotides of the guide to TTCG. For the following positions we reasoned that introducing additional GC biases would destroy 5’-3’ asymmetry of small RNA duplexes that is critically required for efficient RISC loading. To maintain sufficient asymmetry, we therefore decided to bias the next 3 positions towards A or T, which in most positions is in alignment with nucleotide biases associated with knockdown efficacy.

For the remaining sequence 3’ of the seed region, we adhered to nucleotide features associated with knockdown efficacy based on our DSIR analysis, which are remarkably prominent and cannot all be explained through established processing requirements. Altogether, this established a 22-nt matrix for the design of ARTi shRNAmirs (TTCGWWWNNAHHWWCATCCGGN; W = A/T, H = A/T/C; N = A/T/G/C). In a last step, we further reduced possible off-target effects by eliminating all guides whose extended seed sequence (guide positions 2-14) had a perfect match in the human or mouse transcriptome and, finally, selected the following six top-scoring ARTi predictions for experimental validation:

ARTi.6588 – target site: TCCGGATGAAGTTTATATCGAA / shRNAmir (97mer): TGCTGTTGACAGTGAGCGCCCGGATGAAGTTTATATCGAATAGTGAAGCCACAGATGTATTCGATATAAACTTCATCCGGATGCCTACTGCCTCGGA

ARTi.6570 – target site: TCCGGATGATATTGTTATCGAA / shRNAmir (97mer): TGCTGTTGACAGTGAGCGCCCGGATGATATTGTTATCGAATAGTGAAGCCACAGATGTATTCGATAACAATATCATCCGGATGCCTACTGCCTCGGA

ARTi.6634 – target site: TCCGGATGATGTTTTAATCGAA / shRNAmir (97mer): TGCTGTTGACAGTGAGCGCCCGGATGATGTTTTAATCGAATAGTGAAGCCACAGATGTATTCGATTAAAACATCATCCGGATGCCTACTGCCTCGGA

ARTi.6786 – target site: TCCGGATGATATTGTATACGAA / shRNAmir (97mer): TGCTGTTGACAGTGAGCGCCCGGATGATATTGTATACGAATAGTGAAGCCACAGATGTATTCGTATACAATATCATCCGGATGCCTACTGCCTCGGA

ARTi.6834 – target site: TCCGGATGATATTGCATACGAA / shRNAmir (97mer): TGCTGTTGACAGTGAGCGCCCGGATGATATTGCATACGAATAGTGAAGCCACAGATGTATTCGTATGCAATATCATCCGGATGCCTACTGCCTCGGA

ARTi.6516 – target site: TCCGGATGAAGTTTAATTCGAA / shRNAmir (97mer): TGCTGTTGACAGTGAGCGCCCGGATGAAGTTTAATTCGAATAGTGAAGCCACAGATGTATTCGAATTAAACTTCATCCGGATGCCTACTGCCTCGGA

The following ARTi target sequences were used for experimental validation studies: Insertion into the coding sequence before the STOP codon:

ATCCGGATGATATTGTATACGAATCCGGATGATATTGTTATCGAA (with the first “A” being inserted to retain the reading frame)

Insertion after the STOP codon: TCCGGATGATATTGTATACGAATCCGGATGATGTTTTAATCGAATCCGGATGATATTGTTATCGAA shRNAs were ordered as single stranded DNA Ultramer oligonucleotides (Integrated DNA Technologies), amplified by PCR and cloned into different retroviral or lentiviral miRE/miRF shRNAmir expression vectors (LT3GFPIR^6^) using EcoR/XhoI restriction digest or Gibson assembly.

### Cell Culture

Human HCT 116 cells (ATCC: CCL-247) were cultured in RPMI 1640 medium (Thermo Fisher) and PC-9 cells in McCoy’s 5A medium (Thermo Fisher), supplemented with 10% FBS and 1 x GlutaMAX (Thermo Fisher). RKO (ATCC: CRL-2577) and MOLM-13 cells (DSMZ: ACC 584) were cultured in RPMI 1640, supplemented with 10% FBS (Sigma-Aldrich), 4mM L-glutamine (Thermo Fisher), 1mM sodium pyruvate (Sigma-Aldrich), and penicillin/streptomycin (100 U ml^− 1^/100 μg ml^− 1^, Sigma-Aldrich). HT-1080 (ATCC: CCL-121) and LentiX lentiviral packaging cells (Clontech, cat. no. 632180) were cultivated in DMEM (Thermo Fisher) with 10% FBS, 4mM L-glutamine, 1mM sodium pyruvate, and penicillin/streptomycin (100 U ml^− 1^/100 μg ml^− 1^). MV4-11 cells (ATCC: CRL-9591) were cultured in IMDM with 10% FBS, 4mM L-glutamine, 1mM sodium pyruvate, and penicillin/streptomycin (100 U ml^− 1^/100 μg ml^− 1^).

Murine MLL-AF9^OE^, Nras^G12D^ AML cells (RN2; Zuber et al. 2011) were cultured in RPMI 1640 medium supplemented with 10% FBS, 20 mM L-glutamine, 10 mM sodium pyruvate, 10 mM HEPES (pH 7.3), penicillin/streptomycin (100 U ml^− 1^/100 μg ml^− 1^), and 50 μM β-ME. *Kras*^*G12D*^, *Trp53*^-/-^, *MYC*^*OE*^ PDAC cells (EPP2), SV40 large T antigen immortalized mouse embryonic fibroblasts (RRT-MEF^22^; and NIH/3T3 (ATCC: CRL-1658) were cultured in DMEM supplemented with 10% FBS, 20 mM glutamine, 10 mM sodium pyruvate, and penicillin/streptomycin (100 U ml^− 1^/100 μg ml^− 1^). MIA PaCa-2 (ATCC: CRL-1420) and GP2d (Ecacc: 95090714) cells were cultured in DMEM supplemented with 10% FBS. Ba/F3 (DSMZ: ACC300) cells were cultured in RPMI 1640 medium supplemented with 10% FBS, 10ng/ml IL-3 (R&D systems) and Ls513 cells were cultured in RPMI 1640 medium supplemented with 10% FBS. All cell lines were maintained at 37°C with 5% CO_2_, routinely tested for mycoplasma contamination and authenticated by short tandem repeat analysis.

### Reporter Assay

SFFV-GFP-P2A-Puro-ARTi-target sensor was cloned into pRSF91 retroviral plasmid^23^ using Gibson assembly. RRT-MEFs were transduced with retroviruses expressing a GFP-reporter harboring the target sites for validated shRNAs and one ARTi-shRNA in its 3’-UTR. For each reporter cell line single cells were FACS-sorted into 96-well plates using a FACSAria III cell sorter (BD Bioscience) to obtain single-cell derived clones. These clones were transduced with retrovirus constructs in pSin-TRE3G-mCherry-miRE-PGK-Neo (TCmPNe) backbone expressing either the respective doxycycline (dox)-inducible ARTi shRNA or validated shRNAs and mCherry fluorescence marker. shRNA expression was induced with dox and GFP levels were quantified via flow cytometry 2 days post induction. Knockdown efficiency was calculated as 1 minus the ratio of mean GFP signal in mCherry^+^ (shRNA^+^) cells over mCherry^-^ cells and normalized to Renilla luciferase specific neutral control shRNA (Ren.713).

### Competitive proliferation assay

To investigate the effect of ARTi constructs in the absence of the endogenous target gene, competitive proliferation assays were performed in several human and murine cell lines. Human HT-1080, RKO, MOLM-13, and MV4-11 cell lines were lentivirally transduced with shRNAmir expression constructs cloned into pRRL-SFFV-GFP-miRF-PGK-Neo (SGFN) backbone at 20-60% efficiency. Initial infection efficiency was determined at day 4 post transfection (day 0) by measuring GFP expression as a readout using iQue Screener Plus flow cytometer (IntelliCyt). Percentage of shRNA^+^ cells (GFP positive) was monitored by flow cytometry in regular intervals and results were normalized to day 0.

Human GP2d, Ls513, and MIA PaCa-2 cell lines were lentivirally transduced with shRNAmir constructs cloned into pRRL-TRE3G-GFP-miRE-PGK-Puro-IRES-rtTA3 backbone (LT3GEPIR, Addgene plasmid #111177). 500 cells were seeded in duplicates in 96-well plates and treated with 1 µg/ml dox for 9-10 days and analyzed with Incucyte (Sartorius). Untreated cells served as reference.

Murine NIH-3T3, EPP2 and RN2 cells were retrovirally transduced with shRNAmir constructs cloned into pMSCV-miR-E-PGK-Neo-IRES-mCherry backbone (LENC; Addgene plasmid #111163), and initial infection levels were determined by flow cytometry based on mCherry expression 4 days post transduction (day 0).

### Crystal Violet staining

To visualize ARTi-shRNAmir’s effect, crystal violet staining assays were performed. 25’000 HCT 116 cells and 15’000 PC-9 cells per well were seeded in a 6-well plate containing 2 mL tetracycline-free growth medium and 1 µg/mL dox. Medium was exchanged every 2-3 days. After 9 days, wells were washed with ice-cold PBS and subsequently stained with 1 mL of 2.3% crystal violet solution for 10 minutes. Subsequently, wells were washed with ultrapure water and dried overnight. Images were obtained with a scanner.

### Transcriptional profiling

For the unbiased identification of ARTi shRNA off-targets, RKO and HT-1080 shRNA^+^ cells from the competition proliferative assay experiment were selected with Geneticin/G418 Sulfate (Gibco) for 7 days and checked for GFP expression. On day 7, one arm of the empty vector control group was treated with IC_50_ concentration of trametinib (MedChem Express: HY-10999) for 24 hours based on the data in Genomics of Drug Sensitivity in Cancer (GDSC) database (https://www.cancerrxgene.org)^24^. Subsequently, cells were trypsinized, washed with ice-cold PBS, pelleted and snap frozen. Total RNA was isolated using in-house magnetic beads kit and King Fisher Duo Prime Purification System (Thermo Fisher). NGS libraries were prepared with QuantSeq 3’ mRNA-Seq Library Prep Kit (FWD) HT for Illumina (Lexogen) and UMI Second Strand Synthesis Module for QuantSeq FWD (Lexogen). Samples were sequenced on Illumina NovaSeq platform with 100bp single-read protocol.

Engineered MIA PaCa-2 and PC-9 cells were cultured in the presence of 1 µg/ml dox to induce expression of the ARTi shRNA. dox-containing media were replenished twice weekly and on day 4 and day 8 after the initial treatment. 2×10^6^ dox-treated and untreated control cells were harvested, washed with PBS, lysed, treated with DNAse I (QIAGEN) and total RNA was extracted using the RNeasy Mini Kit (QIAGEN). NGS libraries were prepared as above. Samples were sequenced on an Illumina NextSeq 2000 platform with a 75bp protocol.

### Bioinformatic analyses of 3’ mRNA-Seq

For 3’ mRNA-Seq reads derived from the human cell lines HT-1080 and RKO, the six nucleotide long 5’ UMIs were attached to each read name with umi-tools (v1.0.0)^25^. Subsequently, the UMIs plus the next four nucleotides (UMI spacer), as well as 3’ adapters (stringency of 3) and bases with low quality (threshold of 25) were trimmed away using cutadapt (v1.18)^26^ and its wrapper tool trimgalore (v0.6.2). Read quality control was performed with FastQC (v0.11.8). The remaining reads were sample-wise aligned to the human (GRCh38.p13; GCA_000001405.28) reference genome. Mapping and subsequent filtering of 3’ UTR mapped reads was performed with slamdunk (v0.4.3)^27^ in QuantSeq mode (slamdunk map -5 12 -n 100 - q). 3’ UTR regions were assembled based on the description in Muhar et al.^28^. Aligned and filtered reads were deduplicated with umi-tools (v1.0.0) (2), based on the mapping coordinate and the UMI attached to the read name, prior to quantifying read abundances within 3’ UTR regions using featureCounts (v2.0.1)^29^. Differential expression analysis (DEA) was performed with DESeq2 (v1.30.1)^30^ for each ARTi shRNA to empty vector control. Here, the number of up- and downregulated genes were calculated by filtering the DEA results for genes with a log2 fold-change ≥ 2 (up) or ≤ 2 (down) and a -log10 p-value ≥ 5. Principal component analysis was performed on the 1000 most variable expressed genes with the prcomp function from the stats (v4.2.0) R package.

For transcriptional profiling of the human cell lines MIA PaCa-2 and PC-9, the genome reference file and annotations were constructed based on the GRCh38 assembly and the Ensembl 86 version, respectively. The sequences of the shRNA construct (ARTi.6570 (RN_v_76, RN_v_118)) as well as the EGFR^del19^::V5::dsRed::ARTi (RN_v_108) and the dsRed::Linker::KRAS^G12R^-ARTi (RN_v_287) were also included. Mapping of the sequencing reads derived from the human cell lines was performed with STAR (v2.5.2b)^31^ aligner allowing for soft clipping of adapter sequences. Quantification of read counts to transcript annotations was implemented using RSEM (v1.3.0)^32^ and featureCounts (v1.5.1)^29^. Normalization of read counts and differential analysis was implemented with the limma^33^ and voom^34^ R packages.

### Genome engineering of EGFR in PC-9 and Ba/F3

Genome engineering of PC-9 cells was done as previously described^35^. In brief: PC-9_RIEN cells were transduced with an ecotropic pMSCV-EGFRdel19_V5_dsRed_ARTi-PGK-Blasticidin retrovirus cloned at GenScript and produced in Platinum E cells (Cell Biolabs) in the presence of 8 μg/mL Polybrene (Merck Millipore). After 24 hours, stable transgenic cell pools were selected using 10 µg/mL Blasticidin (Sigma-Aldrich). Subsequently, cells were diluted to obtain single cell clones. After 14 days of culture, single cell-derived colonies were transferred to 6-well plates and analyzed by Western Blot. Identified homozygous PC-9_ EGFR^del19^-ARTi clones were further engineered by cutting endogenous EGFR with a CRISPR all-in-one vector pX458_Exon20_gRNA TAGTCCAGGAGGCAGCCGAA (GenScript) using X-tremeGENE 9 DNA transfection reagent (Roche) according to the protocol supplied by the vendor. 48 hours after transfection, GFP positive cells were sorted by FACS (SONY cell sorter S800Z) and diluted to obtain single cell clones. Positive clones, which contained only the exogenous EGFR^del19^-ARTi, but not the endogenous EGFR, were identified by Western Blot. Next, the selected EGFR clone was transduced with a pantropic LT3GEPIR_Puro_ARTi-shRNA TTCGATAACAATATCATCCGGA retrovirus cloned at GenScript, China and produced via the Lenti-X Single Shot system (Clontech). 72 hours later, stable transgenic cell pools were selected using 0.5µg/mL Puromycin (Sigma-Aldrich). Following the selection, cells were diluted to obtain single cell clones. After 14 days of culture, single cell-derived colonies were transferred to 6-well plates and induced via 1µg/ mL dox. Positive clones were characterized by a strong GFP-induction that was identified by flow cytometry.

Ba/F3 cells were transduced with an ecotropic pMSCV-EGFRdel19_V5_dsRed_ARTi-PGK-Blasticidin retrovirus cloned at GenScript and produced in Platinum E cells in the presence of 4 μg/mL Polybrene. After 72 hours, stable transgenic cells were selected by using 50 µg/mL Blasticidin, without adding IL-3.

### Genome engineering of KRAS in MIA PaCa-2

MIA PaCa-2 ARTi-shRNAmir expressing cells were transduced with an ecotropic pMSCV-dsRed::KRAS^G12R^-ARTi-PGK-Blasticidin retrovirus cloned at GenScript, China and produced in Platinum E cells in the presence of 8 μg/mL Polybrene. After 24 hours, stable transgenic cell pools were selected using 10 µg/mL Blasticidin. Subsequently, endogenous KRAS was knocked out by transient transfection of three gRNAs targeting exon 2 and the region containing the G12C variant (present in MIA PaCa-2 cells) were used in a co-transfection (gRNA#3: *GAATATAAACTTGTGGTAGT*; gRNA#6: *CTTGTGGTAGTTGGACTTG*; gRNA#7: *GTAGTTGGAGCTTGTGGCGT*). Knockout clones were identified by the absence of the endogenous KRAS protein using Western Blot.

### Genome engineering of STAG1/STAG2 in HCT 116

HCT 116 cell line was engineered by cutting STAG1 with gRNAs targeting the region close to the STOP codon. Guide RNAs *TTCTTCAGACTTCAGAACAT* or *CTGAAGAAAATTTACAAATC* were cloned into the pSpCas9(BB)-2A-GFP plasmid (pX458; Addgene plasmid 48138) and used in a co-transfection. Simultaneously, a STAG1_AID_V5_P2A_Blasti_STOP_ARTi repair template with 800bp of left and right homologous arms of the STAG1 genomic locus (in pUC57-Simple backbone) was transfected into HCT 116 cells using Lipofectamine 3000 transfection reagent (Thermo Fisher) according to the manufacturer’s instructions. A stable transgenic cell pool was selected 48 hours after transfection using 5µg/ml Blasticidin and diluted to obtain single cell clones. Positive clones were identified by Western Blot.

Identified homozygous HCT 116_STAG1_ARTi clones were further engineered by disrupting STAG2 gene with a CRISPR all-in-one vector Hs0000077505_U6gRNA-Cas9-2A-GFP and Hs0000077502_U6gRNA-Cas9-2A-GFP, respectively. Cells were transfected using X-tremeGENE 9 DNA transfection reagent according to the manufacturer’s instructions, sorted 48 hours post transfection for GFP positive cells and diluted to obtain single cell clones. Positive clones were identified by Western Blot.

Next, the selected HCT 116 clone was transduced with a pantropic LT3GEPIR_Puro_ARTi-shRNA *TTCGATAACAATATCATCCGGA* retrovirus cloned at GenScript, China and produced via the Lenti-X Single Shot system (Clontech). 72 hours later, stable transgenic cell pools were selected using 2 µg/mL Puromycin (SIGMA, P9620). Following the selection, cells were diluted to obtain single cell clones. After 14 days of culture, single cell-derived colonies were transferred to 6-well plates and induced via 1µg/ mL dox. Positive clones were characterized by a strong GFP-induction that was identified by flow cytometry.

### Western Blot

The following primary antibodies were used for immunoblot analyses: EGFR (Cell Signaling, #4267); STAG1 (GeneTex, GTX129912); STAG2 (Bethyl, A300-159A); b-actin (Sigma, A5441); KRAS (LSbio, LS-C17566); V5 (Sigma, V8012). PC-9, MIA PaCa-2, and HCT 116 cell pellets harboring EGFR, KRAS, and STAG1/STAG constructs respectively were lysed in Triton X-100 lysis buffer, sonicated and stored at -80°C. For protein detection, the pellets were thawed on ice, followed by 15 minutes centrifugation at 13,000 rpm and 4°C. Further, cell lysates were loaded onto a pre-casted SDS–polyacrylamide gel (4-12%) and proteins were transferred onto a nitrocellulose or PVDF-membrane. Membranes were probed with the respective primary antibodies overnight. The next day, secondary antibodies conjugated with fluorescent dye were added and the proteins were detected by the Odyssey detection system.

### Compound treatment

To investigate sensitivity to EGFR-targeting compounds, cell viability was determined using the Cell Titer Glo assay (Promega). For this purpose, 10 mM stock solutions in DMSO of afatinib^36^, Osimertinib^37,38 37^ and poziotinib^39,40^ were used. 5’000 cells per well were seeded in 150µL of the medium in technical triplicates in 96-well plates and incubated at 37°C and 5% CO_2_ for 5 hours, followed by the compound addition. Cells were treated with seven different concentrations of inhibitors in a serial eight-fold dilutions starting with the highest concentration of 3 µM. For comparability, DMSO normalization to the highest added volume was performed. Subsequently, cells were cultivated for 96 hours at 37°C and 5% CO_2_. 50 µL of Cell Titer Glo reagent was added to each well, incubated for 10 minutes in the dark and luminescence was measured using the multilabel Plate Reader VICTOR X4. The measurement time was set to 0.2 seconds. Luminescence values relative to DMSO-treated cells were plotted in GraphPad Prism and fitted using nonlinear regression with a variable slope to calculate IC50 values at 50% inhibition. MIA PaCa-2 KRAS G12C inhibitor (AMG-510)^17–19^ treatments were performed as described for EGFR with the following modifications: 2’000 cells per well, and 0.5nM-3µM concentration range of AMG-510.

### *In vivo* experiments

The PC-9 EGFR k.o., EGFR^del19^::ARTi study was performed at the AAALAC accredited animal facility of CrownBio Leicestershire, UK. Female NSG® (NOD.Cg-PrkdcscidII2rgtm1wjl/SzJ) Crl mice were obtained from Charles River. Age of animals at study initiation was 7-8 weeks and hat an acclimatization period of ≥14 days. Mice were groups housed in IVCs. The study complies with the UK Animals Scientific Procedures Act 1986 (ASPA) in line with Directive 2010/63/EU of the European Parliament and the Council of 22 September 2010 on the protection of animals used for scientific purposes.

All other in vivo experiments were performed at the AAALAC accredited animal facility of Boehringer Ingelheim RCV GmbH & CoKG. Female BomTac:NMRI-*Foxn1*^*nu*^ mice were obtained from Taconic Denmark at 6-8 weeks of age. After the arrival, mice were allowed to adjust to the housing conditions at least for 5 days before the start of the experiment. Mice were housed in pathogen-free and controlled environmental conditions (open cage housing), and handled according to the institutional, governmental and European Union guidelines (Austrian Animal Protection Laws, GV-SOLAS and FELASA guidelines).

Studies were approved by the internal ethics committee (called “ethics committee”) of Boehringer Ingelheim RCV GmbH & Co KG in the department of Cancer Pharmacology and Disease Positioning. Furthermore, all protocols were approved by the Austrian governmental committee (MA 60 Veterinary office; approval numbers GZ: 903122/2017/21 and GZ: 416181-2020-29).

To establish subcutaneous tumors, mice were injected with 2 × 10^6^ HCT 116 in PBS, 5 × 10^6^ MIA PaCa-2 in 1:2 Matrigel : PBS with 5% FBS or with 1 × 10^7^ PC-9 cells in PBS with 5% FBS.

Tumor diameters were measured with a caliper three times a week. The volume of each tumor [in mm^3^] was calculated according to the formula “tumor volume = length * diameter^2^ * π/6.” Mice were randomized into the treatment groups when tumor size reached between ∼130 and 190 mm^3^. Group sizes were calculated for each tumor model based on tumor growth during model establishment experiments. A power analysis was performed using a sample size calculator (https://www.stat.ubc.ca/~rollin/stats/ssize/n2.html). For all models used in the studies, 10 mice per group were used. 2mg/ml doxycycline hyclate (Sigma) and 5mg/ml sucrose were added to the drinking water of the treatment groups, the control group received water with 5mg/ml sucrose only. Osimertinib (Tagrisso, AstraZeneca, 40 mg tablet) was dosed per os daily at a dose of 25mg/kg in Natrosol and control mice were dosed per os daily with Natrosol. To monitor side effects of treatment, mice were inspected daily for abnormalities and body weight was determined three times per week. Animals were sacrificed when the tumors reached a size of 1’500 mm^3^. Food and water were provided ad libitum. *In vivo* experiments were not repeated.

### Immunohistochemistry

Xenograft samples were fixed in 4% formaldehyde for 24 h and embedded in paraffin. Two-micrometer-thick sections were cut using a microtome. STAG1 was stained with Polink 2 Plus HRP rat NM detection system (GBI Labs #D46-6) according to manufacturer’s instructions using a recombinant rat monoclonal STAG1 antibody (Abcam ab241544, Lot:GR3334172-1; 1:200) after cooking 10 minutes at 110°C in antigen unmasking solution (Vector #H3301). After staining the slides were digitalized (scanner: Leica Aperio AT2). All slides were reviewed and evaluated for quality by a board-certified MD Pathologist.

Imaging analysis was performed using the digital pathology platform HALO (Indica Labs). A tissue-classifying algorithm was trained to selectively recognize viable tumor tissue against stroma, necrosis, and skin. The tissue classification output for each scan was reviewed and manually edited as necessary. A cell detection and scoring algorithm was trained to measure DAB optical density (OD) in the nuclei of tumor cells. A positivity threshold for DAB OD was determined by normalization with respect to the DAB OD as calculated from *bona fide* negative tissue (e.g., murine stroma as background). The object data for background-normalized nuclear ODs for each tumor cell were exported from three control and three doxycycline-treated cases (207945 and 33331 pooled objects, respectively). The cumulative distribution of the background-normalized DAB OD for each tumor cell nucleus was then plotted for control and doxycycline-treated cases, separately (supplemental figure 4, panel K).

### Reagents, code and data availability

DNA sequences of the transgenes used in this study and the code used to generate the figures are available upon request. The workflow for the RNA-seq bioinformatics analyses is described in the Materials and Methods section. RNA sequencing data were uploaded to GEO with the accession numbers: GSE218404 and GSE218617. The data will be made publicly available upon acceptance of this manuscript. All ARTi cell lines and constructs described in this study are available upon request.

### Cell lines

All newly generated ARTi cell lines are available upon request. The following parental cell lines were used:

| Cell line name | species | Supplier name | Catalog number | Clone number |
| --- | --- | --- | --- | --- |
| PC-9 | human | ECACC | 90071810 | Lot# 14A030 |
| MIA PaCa-2 | human | ATCC | CRL-1420 | Lot# 1350798 |
| HT 1080 | human | ATCC | CCL-121 | Lot# 63835574 |
| HCT 116 | human | ATCC | CCL-247 | Lot# 6028631 |
| RKO | human | ATCC | CRL-2577 | Lot# 59354083 |
| MV4-11 | human | ATCC | CRL-9591 (old number) | Lot# 58352230 |
| MOLM-13 | human | DSMZ | ACC 554 | Lot# 11 |
| NIH-3T3 | human | ATCC | CRL-1658 | Lot# 63292276 |
| EPP2 |  | KrasG12D, Trp53<sup>-/-</sup>, MYCOE PDAC cells (EPP2) | NA | NA |
| RN2 | murine | MLL-AF9OE, NrasG12D AML cells (RN2; Zuber et al. 2011) | NA | NA |
| Gp2d | human | ECACC | 95090714 | Lot# 11E013 |
| Ls513 | human | ATCC | CRL-2134 | Lot# 57761342 |
| Ba/F3 | murine | DSMZ | ACC300 |  |
| 293T-Lenti-X | human | Clontech | 632180 | Lot# 1404558A |

## Acknowledgements

The authors thank all colleagues in the labs of Johannes Z. and R.A.N. for critical discussions and technical support.

