## Supplementary figures and images for "Precision RNAi using synthetic shRNAmir target sites"

### Supplemental Data 1

#### Design and selection of ARTi-shRNAmirs

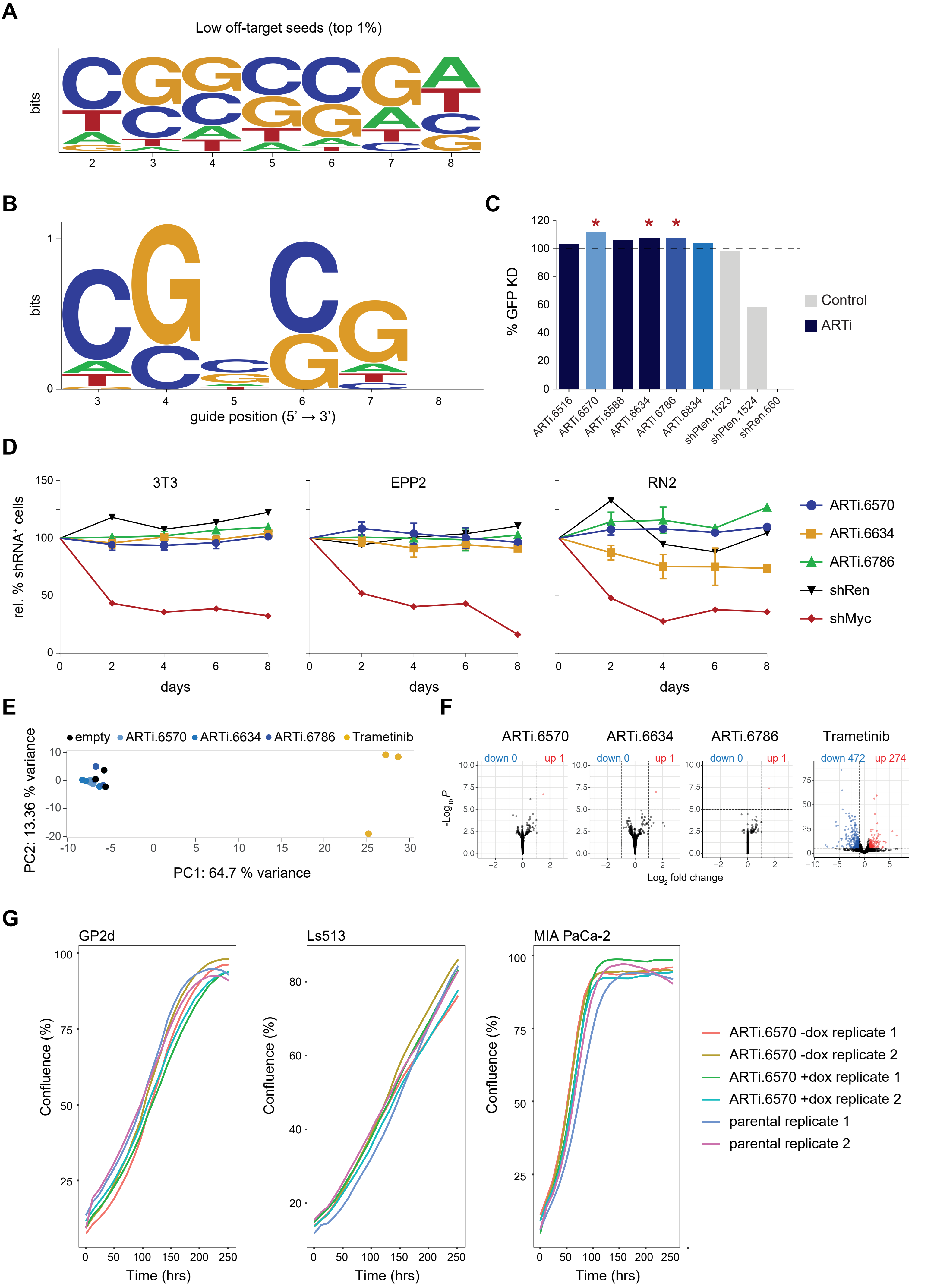


Hoffmann et al., Figure 1 – figure supplement 1

### Supplemental Data 4

#### Validation of ARTi in vitro and in vivo - STAG1

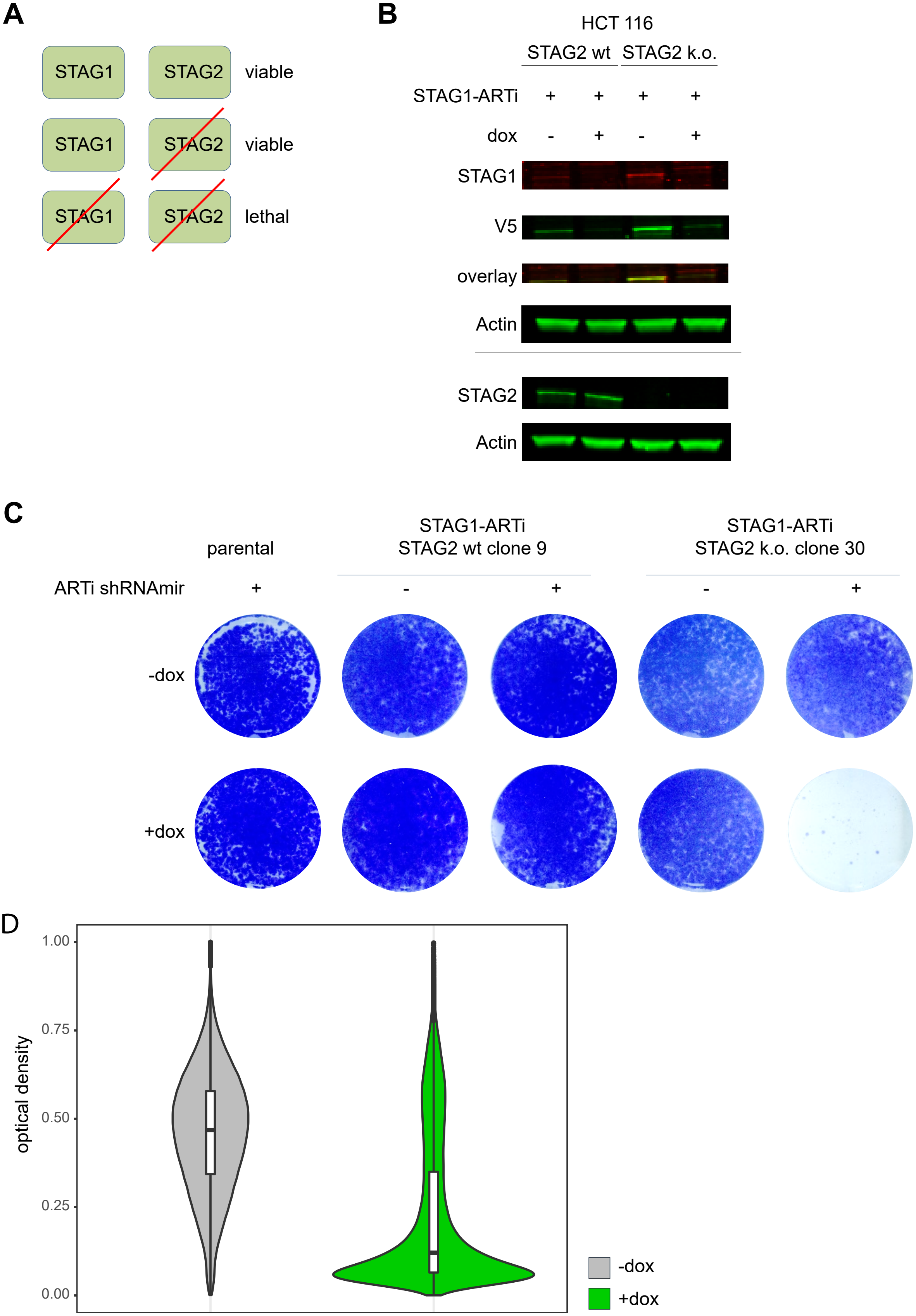


Hoffmann et al., Figure 2 – figure supplement 3
