## Supplemental Data 2 for "Precision RNAi using synthetic shRNAmir target sites"

### Validation of ARTi in vitro and in vivo - EGFR

**A**

| Ba/F3 cells $GI_{50}$ [nM] | IL-3 (parental) | $EGFR^{del19}$ | $EGFR^{del19,C797S}$ | $EGFR^{del19}$ ARTi |
|---|---|---|---|---|
| Afatinib | >500 | 0.14 | 3.3 | 0.06 |
| Poziotinib | >500 | 0.06 | 80 | 0.03 |
| Osimertinib | >2000 | 0.03 | >2000 | 3.6 |

**B**

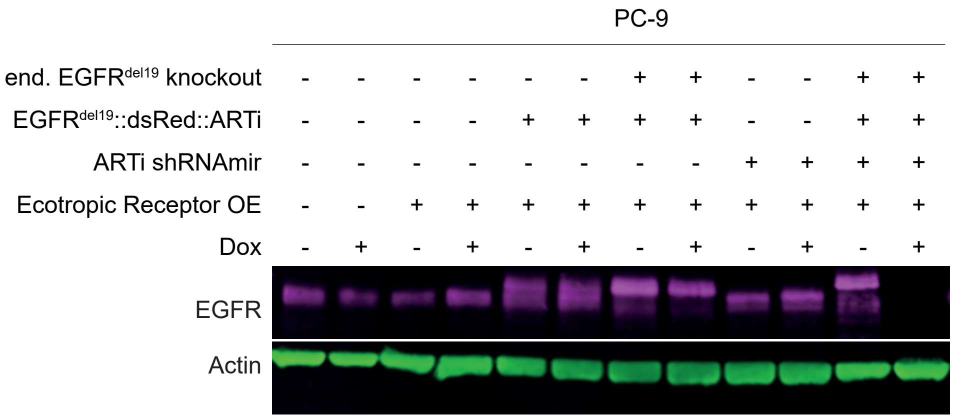


**C**

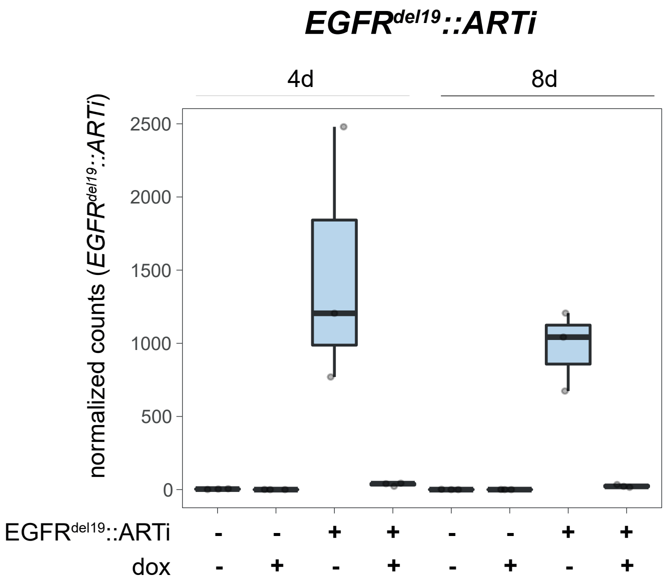


**D**

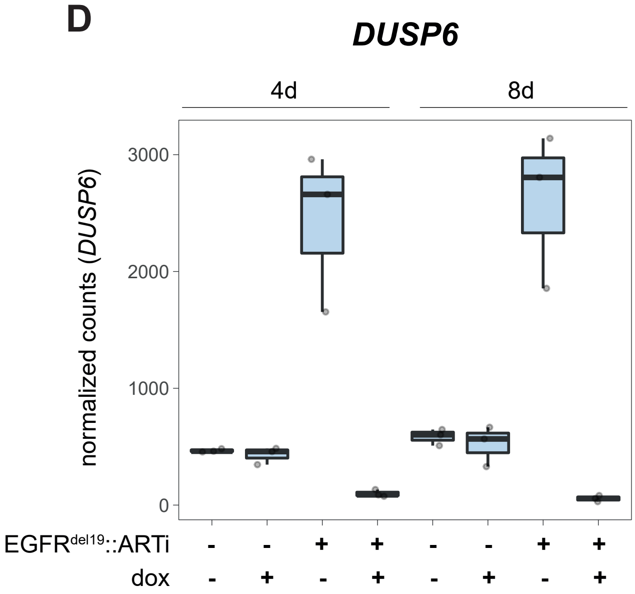


**E**

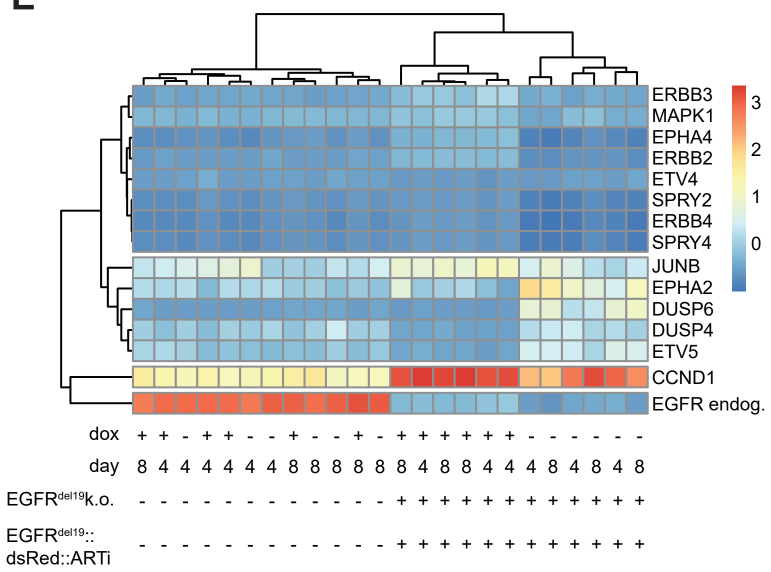


**F**

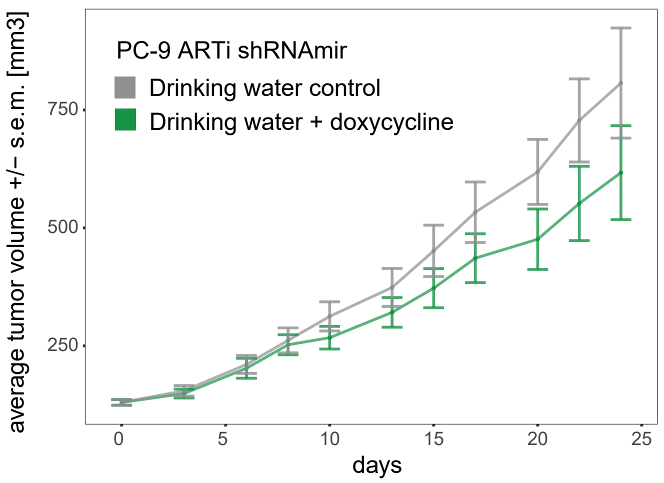
