## Supplemental Data 3 for "Precision RNAi using synthetic shRNAmir target sites"

**Validation of ARTi in vitro and in vivo - KRAS**

**A**

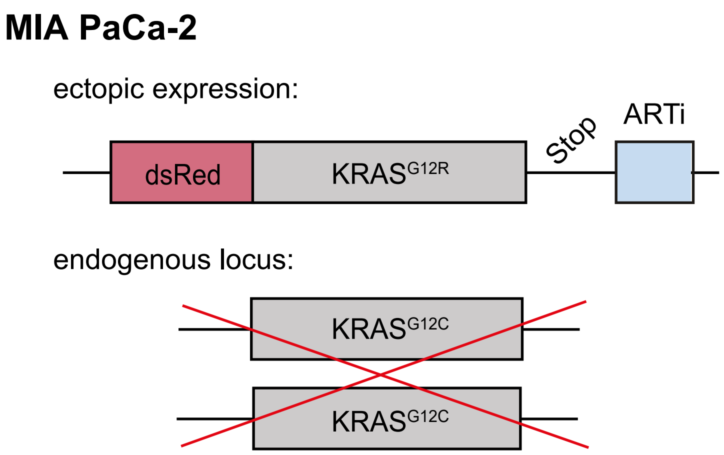


**B**

| KRAS$^{G12C}$i (AMG-510) | $GI_{50}$ [nM] |
|---|---|
| MIA PaCa-2 parental | 52 |
| MIA PaCa-2 KRAS$^{G12C}$ k.o. dsRed::KRAS$^{G12R}$-ARTi ARTi-shRNAmiR (-dox) | >5000 |

**C**

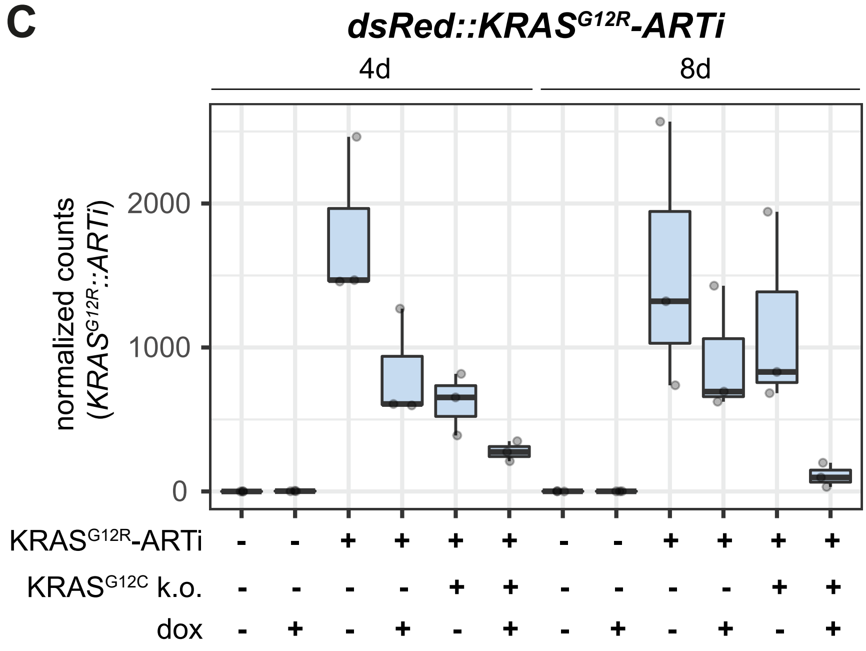


**D**

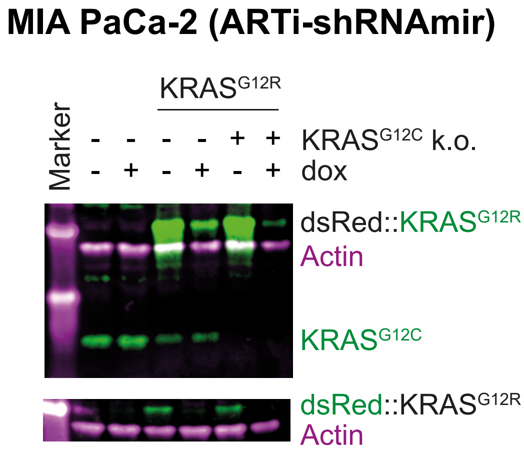
